# Identifying disease-associated circRNAs based on edge-weighted graph attention and heterogeneous graph neural network

**DOI:** 10.1101/2022.05.04.490565

**Authors:** Chengqian Lu, Lishen Zhang, Min Zeng, Wei Lan, Jianxin Wang

## Abstract

**Motivation:** Circular RNAs (circRNAs) with varied biological activities are implicated in pathogenic processes, according to new findings. They are regarded as promising biomarkers for the diagnosis and prognosis due to their structural features. Computational approaches, as opposed to traditional experiments, can identify the circRNA-disease connections at a lower cost. Multi-source pathogenesis data can help to reduce data sparsity and infer probable connections at the system level. The majority of available approaches create a homologous network using multi-source data, but they lose the data’s heterogeneity. Effective solutions that make use of the peculiarities of multi-source data are urgently needed.

**Results:** In this paper, we propose a model (CDHGNN) based on edge-weighted graph attention and heterogeneous graph neural networks for discovering probable circRNA-disease correlations prediction. The circRNA network, miRNA network, disease network and heterogeneous network are constructed based on the introduced multi-source data on circRNAs, miRNAs, and diseases. The features for each type of node in the network are then extracted using a designed edge-weighted graph attention network model. Using the revised node features, we learn meta-path contextual information and use heterogeneous neural networks to assign attention weights to different types of edges. CDHGNN outperforms state-of-the-art algorithms with comparable accuracy, according to the findings of the trial. Edge-weighted graph attention networks and heterogeneous graph networks have both improved performance significantly. Furthermore, case studies suggest that CDHGNN is capable of identifying particular molecular connections and can be used to investigate pathogenic pathways.

## 1 Introduction

Back-spliced from precursor mRNAs, circular RNA (circRNA) is a single-stranded endogenous non-coding RNA with covalently closed loop structures (Jeck *et al*., 2014). Emerging evidences show that circRNAs play a variety of functions, such as transcriptional regulators, microRNA sponges and protein templates (Huang *et al*., 2020). For example, exon 15 of SMARCA5 is stopped because that circSMARCA5 becomes an R-loop after binding to its parent gene locus (Xu *et al*., 2020). As a consequence, abnormal expression or dysfunction of circRNA will rise a variety of diseases. Compared with linear transcripts, circRNA is more stable without free ends that are susceptible to exonuclease digestion. Therefore, circRNA is expected to be a promising diagnostic biomarker due to its unique structure and biological roles. Nonetheless, identifying experimentally verified disease-related circRNAs takes a lot of manpower and material resources.

Computational methods provide effective means for large-scale discovery of circRNA-disease associations. Existing approaches are categorized into three types. The first type of methods are based on network propagation algorithms. The KATZ measure is used to calculate the potential association probabilities based on a heterogeneous network formed from various biological data. (Fan *et al*., 2018; Zhao *et al*., 2019; Deng *et al*., 2019). Lei *et al*. (2020) proposed a model to classify associations based on a heterogeneous network with the random walk with start. Only topological information is taken into account, limiting the model’s learning ability.

The second type of methods are based on machine learning models. Yan *et al*. (2018) presented a method based on the regularized least-squares of Kronecker product kernel with disease semantic similarity and Gaussian interaction profiles. Wei *et al*. (2020) applied matrix factorization based on disease semantic information, circRNA-gene, gene-disease and circRNA-disease. Xiao *et al*. (2019) employed a weighted low-rank approximation model with dual-manifold regularization on Gaussian interaction profile kernel, disease network and circRNA network. Wei *et al*. (2021) proposed a ranking model to obtain global ranking relationships between query circRNAs and diseases. Based on low-dimensional node representation, Xiao *et al*. (2021) proposed a network embedding-based adaptive subspace learning method to infer potential associations. The appropriate features necessitate professional knowledge and have an impact on classifier performance. The third type of methods are based on neural networks. Lu *et al*. (2020) applied neural networks to replace linear approximation based on matrix factorization for possible associations. Wang *et al*. (2020) extracted features of circRNAs and disease from multi-source and classified associations with ELM classifier. Lu *et al*. (2020) employed an unsupervised model to learn *k*-mer low-dimensional vectors and disease ontology representation, as well as BiLSTM to obtain circRNA-disease connections. Wang *et al*. (2020) gained features from biological data with stacked autoencoder algorithm and predicted associations with rotation forest classifier. After fusing features of circRNA sequence and disease semantics, Wang *et al*. (2020) pre-trained the generative adversarial networks with circRNA-disease pairs and inferred prediction with the extreme learning machine classifier. Ignoring the close connection between features of circRNAs (diseases) and their associations prevents the models from learning intrinsic representation. Mudiyanselage *et al*. (2021) applied graph convolutional networks on the constructed heterogeneous network. Lan *et al*. (2021) integrated multiple relationship among circRNA, miRNA, lncRNA and disease based on graph attention network. The methods mentioned above have improved accuracy of prediction. However, the heterogeneity between different data is disregarded when applying graph neural networks on multi-source data.

In this paper, we offer a heterogeneous graph neural network model (named CDHGNN) that can learn not only the hidden properties of each type of biological molecule, but also the heterogeneity between different source data. We introduce multi-source data of circRNA, miRNA and disease to alleviate data sparsity and explore molecular associations. After that, we construct circRNA network, miRNA network, disease network and heterogeneous network. We devise an edge-weighted graph attention network to get node representation, in contrast to prior graph neural network methods that disregard edge weights. Furthermore, we adopt heterogeneous transformer network to learn the contextual information of the meta-path and get attention weights for different types of edges. The experimental results reveal that CDHGNN outpaces state-of-the-art computational methods. Edge-weighted graph attention network and heterogeneous graph network improve accuracy. It is worth noting that CDHGNN finds molecular connections and the relevant pathways in pathogenesis.

The contributions of this work are summarised as follows:

- To view molecular associations in pathogenesis, we integrate multi-source data of circRNA, miRNA and disease and construct corresponding biological networks.
- We devise an edge-weighted graph attention neural network that considers the value of associations when obtaining node intrinsic properties.
- We first study the heterogeneity of multi-source data. CDHGNN learns the meta-contextual path’s information and assigns attention weights to various types of edges based on a heterogeneous neural network.
- The experimental results and case studies show that CDHGNN out-performs state-of-the-art methods and is helpful to explore relevant pathways in pathogenesis.

## 2 Materials and Methods

In this part, we present our model CDHGCN, which is applied to identify the potential circRNA-disease associations. As shown in Fig. 1, we first construct circRNA network, miRNA network, disease network and heterogeneous network. Then, we extract node features with a devised edge-weighted graph attention network. Finally, we learn contextual information and assign attention weights on the meta-path based on a heterogeneous neural network.

**Fig. 1.**
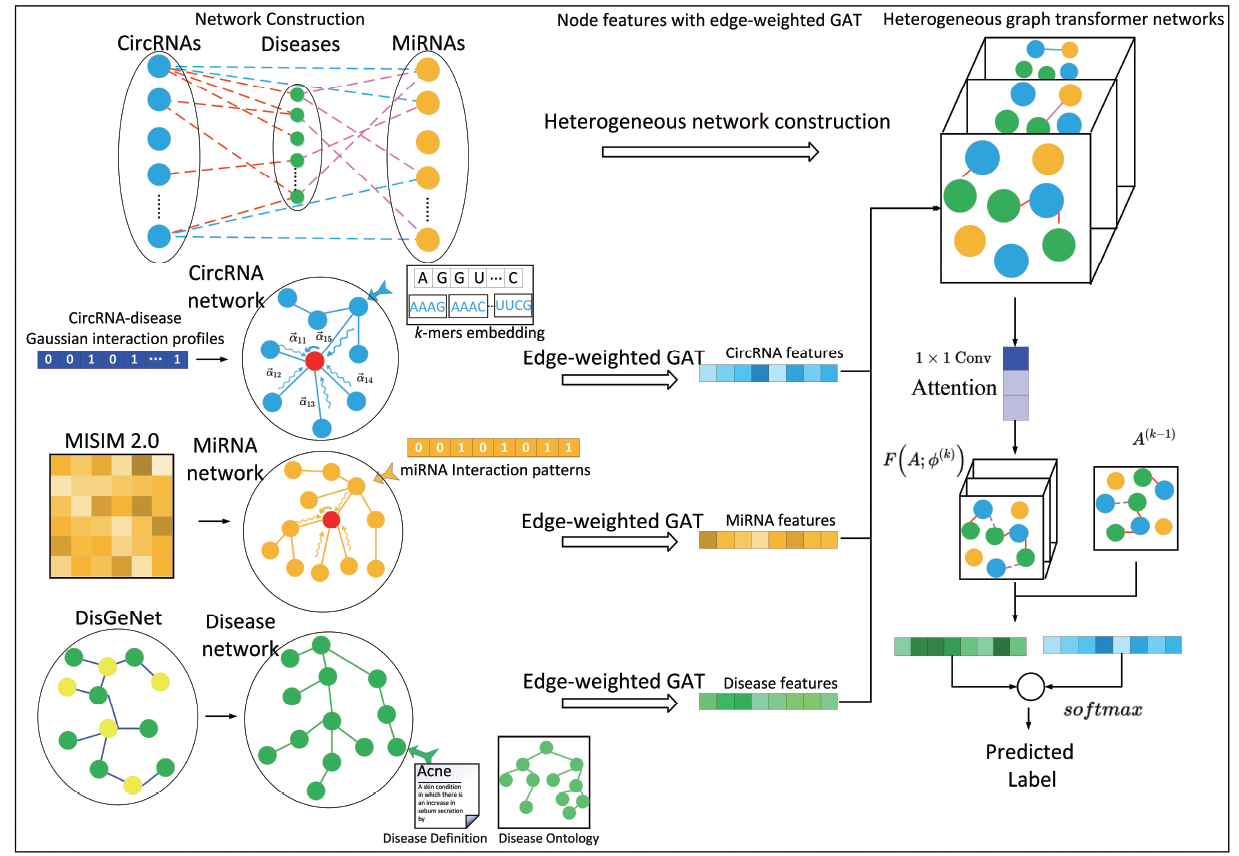
The overflow of CDHGNN. Step 1: construct circRNA-circRNA network, miRNA-miRNA network, disease-disease network and heterogeneous network; Step2 : extract each type of node features with edge-weighted graph attention network; Step3: Learn contextual information and assign attention weights on the meta-path in the heterogeneous network.

### 2.1 Benchmark Dataset

To alleviate data sparsity and understand potential associations systematically, we collect circRNA sequences, miRNA functional relationships, disease ontology, gene-disease network, circRNA-miRNA, miRNA-disease and circRNA-disease associations from benchmark database. We retrieve circRNA sequences data from CircBase (Glazar *et al*., 2014) and get 140,732 circRNA sequences. We download miRNA functional relationships from database MISIM v2.0 (Li *et al*., 2019), which includes functional relationships among 664 miRNAs. We obtain disease ontology from Disease Ontology (Schriml *et al*., 2019), which contains 11,652 phenotype ontology. The UMLS Metathesaurus Browser, a vast biomedical thesaurus, is used to get disease definitions. We get gene-disease associations from (Pinero *et al*., 2020), which contains 262,989 associations between 13,705 genes and 1,977 diseases. We gather circRNA-miRNA associations from starBase (Li *et al*., 2014), which contains 18,320 circRNA-miRNA associations between 886 circRNAs and 638 miRNAs. We get miRNA-disease associations from HMDD 3.0 (Huang *et al*., 2019), which includes 27,872 miRNA-disease associations between 1,054 miRNAs and 226 diseases. We download circRNA-disease associations from MNDR 3.0 (Ning *et al*., 2021), which includes 3,206 circRNA-disease associations between 2,396 circRNAs and 165 diseases. We have unified and standardized the nomenclature of circRNA according to circBase. The unified and standardized operation is executed for the nomenclature of disease according to OMIM and UMLS. After that, we delete duplicate data of species other than human species. Finally, we get 2,013 associations between 1,313 circRNAs and 144 diseases, 10,570 associations between 638 miRNAs and 128 diseases, and 13,315 associations between 824 circRNAs and 612 miRNAs. The details of the data are depicted in Table 1.

**Table 1.** The details of the data

| Dataset | Num |
| --- | --- |
| Disease ontology | 11,652 |
| Disease definition | 152 |
| CircRNA sequences | 140,732 |
| MiRNA-disease | 10,570 (638 miRNAs, 128 diseases) |
| CircRNA-miRNA | 13,315 (824 circRNAs, 612 miRNAs) |
| CircRNA-disease | 2,013 (1313 circRNAs, 144 diseases) |

### 2.2 Network construction

After importing multi-source data on the pathogenesis of circRNAs, heterogeneous biological network is an effective method to find the promising associations between circRNAs and diseases. There are three types of nodes like circRNA, miRNA and disease. To obtain features of each type of node in the respective biological network, CDHGNN establishes circRNA network, miRNA network and disease network, respectively.

The Gaussian interaction profiles (GIPs) embody association patterns of circRNA or disease (Lu *et al*., 2018). We get the association matrix between circRNAs and diseases *A*_*cd*_ ∈ ℝ^*nc*×*nd*^, where *nc* is the number of circRNAs while *nd* is the number of diseases, respectively. We calculate the Gaussian interaction profile kernel similarity as the similarity between two circRNAs:

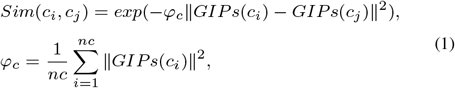

where *φ*_*c*_ controls the kernel bandwidth.

To build a miRNA functional similarity network, we download functional relationships of miRNA from MISIM v2.0. To make the values have the same scale, we normalized the data with Z-score normalization and use it as the functional similarity among miRNAs.

To construct disease network, we download gene-disease data from DisGeNET. We get the association probability between genes and diseases. Then, we calculate disease similarity between disease *i* and disease *j* as follows:

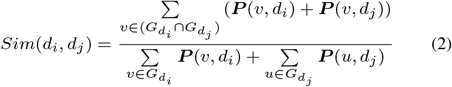

where 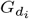 is the set of genes related with disease 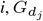 is the set of genes related with disease j, where P(·, ·) is the association probability matrix between genes and diseases.

There are three type of edges among circRNAs, miRNAs and disease, such as circRNA-miRNA, circRNA-disease and miRNA-disease. A heterogeneous network is defined as *G* = (*V, E*), where *V* and *E* signify the sets of nodes and edges, respectively. The heterogenous network is accompanied by a node type mapping function *ψ*_*v*_ : *V* → 𝒯^*v*^ and an edge type mapping function *ψ*_*e*_ : *E* → 𝒯^*e*^. A node type mapping *ψ*_*v*_*(v*_*i*_) uniquely corresponds to a node *v*_*i*_, *i*.*e*., *ψ*_*v*_(*v*_*i*_) ∈ 𝒯^*v*^, *v*_*i*_ ∈ *V*. Analogously, an edge type mapping *ψ*_*e*_(*e*_*i*_) uniquely corresponds to an edge *e*_*i*_, i.e., *ψ*_*e*_(*e*_*i*_) ∈ 𝒯^*e*^, *e*_*i*_ ∈ *E*. The situation is |𝒯_*e*_| *>* 1 in the heterogeneous network. We define the heterogeneous network with a set of adjacency matrices 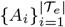, where *A*_*i*_ ∈ ℝ^*N* × *N*^, *N* = *nc* + *nm* + *nd*. A node feature matrix *X* ∈ ℝ^*N* × *F*^ is the input vector to the heterogeneous network, where F denotes the learned features for N nodes from GAT. A meta-path *P* is a path through the heterogeneous network, i.e., 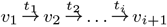, where *t*_*i*_ ∈ 𝒯^*e*.^ Then the the adjacency matrix *A*_𝒫_ of the meta-path 𝒫 is defined as 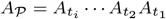, where 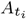 is an adjacency matrix for the *i*-th type of edges on the meta-path.

### 2.3 Node embeddings with edge-weighted graph attention networks

We extract features of various types of nodes from the three constructed similarity networks. As each molecule functions differently in biological systems, we apply graph attention network (GATs) to specify different weights to different nodes in a neighborhood (Velickovic *et al*., 2018). To account for the effect of edge weights on aggregation, we devise a novel edge-weighted GATs. Input a set of initial node features, 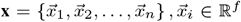, where *n* means the number of nodes, and *f* is the feature dimension of each node. Transform the initial features into high-level features with a linear operation 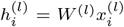, where *W* is a shared weight matrix and 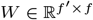. The generated features of *l*-layer is 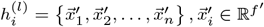. The edge weight between node *I* and node *j* reflects the importance on each connection. Concatenate 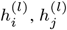 and edge weight *ω*_*ij*_ between node *i* and node *j* in *l* l-th layer and calculate a un-normalized pair-wise attention score as:

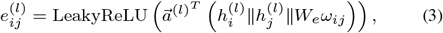

where 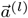 is a learnable weight vector, 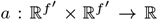. Masked attention is applied to insert graph structure into the model architecture.

Normalizing coefficients across all choices of j with the softmax function:

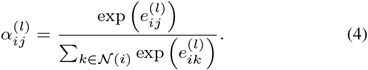

Multi-head attention aggregation is adopted to aggregate the embeddings of neighbors together simultaneously

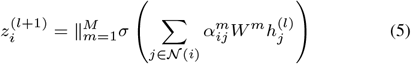

For the initial node features in circRNA network, CDHGNN utilizes Doc2Vec algorithm (Le *et al*., 2014) to get sequence motif embedding on the seuqnce. For the initial node features in miRNA network, miRNAs’ GIPs, 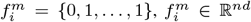, are employed as association vectors. For disease, we concatenate disease semantic similarity based on disease ontology and disease semantic embedding from disease definitions with Doc2Vec.

### 2.4 Heterogeneous graph transformer networks

Heterogeneous networks composed of multiple types of data are models successfully applied in various applications. However, most GNN models treat this type of complex network including multiple types of data as a homogeneous network, or need to manually specify meta-paths in advance. These methods may cause loss of information. In this part, we applied graph transformer networks (GTN) to learn soft selections of edge types and get hidden relationships among circRNAs, miRNAs and diseases (Yun *et al*., 2019). The input to GTN is multiple networks with diverse types of nodes and edges. The 1×1 convolution layers assigns a weight to each type of edge in 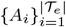, and then softmax(*W*_*ϕ*_) utilizes attention mechanism to determine the influence on the final meta-path. A soft adjacency matrix is generated from the weighted sum defined as:

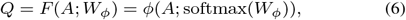

where *φ* is the convolution layer and 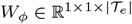. Then, a composition meta-path is established by multiplying the adjacency matrices. To consider the characteristics of the original edges, we introduce the identity matrix *I*. Applying GCN for each type of edge on the meta-path, node representations are shaped as:

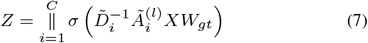

where ‖ denotes the concatenate operator, C is the number of channels, 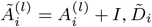 is the degree matrix of 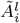, and *W*_*gt*_ ∈ ℝ^*F* × *F*^ is a shared trainable weight matrix. We utilize the cross-entropy loss function to measure the performance and the Adam algorithm to optimize the model.

## 3 Results and Discussion

### 3.1 Evaluation metrics

We implement 5-fold cross-validation to verify the effectiveness of the model and compare it with state-of-the-art methods. All the verified associations among circRNA, miRNA, and disease are treated as positive samples, while candidate samples are potential relationships that have not been validated. To alleviate the imbalance, we stochastically generate negative samples with the same number of positive samples. Then all the samples are randomly divided into 5 parts. One of them is treated as a separate test set that does not participate in the model’s training. The remaining 4 parts take part in the training. We repeat the whole process five times until each part is tested once. THe general evaluation criteria are utilized, including Accuracy (ACC), Precision (Pre), Recall and F1-score defined as follows:

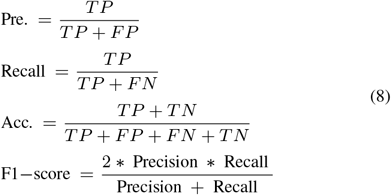

where TP and TN are the numbers of correctly identified positive and negative samples, FP and FN are the number of incorrectly identified positive and negative samples, respectively.

### 3.2 Effects of Parameters

In this part, we evaluate the impact of several key parameters on the model, including learning rate and node dimension. The learning rate is a critical parameter for the minimum of the loss function. We execute the grid search to obtain the optimal value from 0.001 to 0.3. From Table 2, it shows that CDHGNN performs the best with a value of 0.005. As the parameter increases from 0.001 to 0.005, the accuracy of the model is improving. But as the parameter continues to increase from 0.005, the performance of the model has declined. A large learning rate will cause the model to converge to a sub-optimal solution. The default value of the learning rate is set to 0.005.

**Table 2.** The impact of learning rate on model performance

| Learning rate | AUC | AUPR | Acc | Pre | Recall | F1-Score |
| --- | --- | --- | --- | --- | --- | --- |
| 0.001 | 0.851 | 0.792 | 0.817 | 0.799 | 0.786 | 0.792 |
| 0.003 | 0.878 | 0.801 | 0.822 | 0.782 | 0.792 | 0.803 |
| **0.005** | **0.886** | **0.817** | **0.824** | **0.808** | **0.817** | **0.804** |
| 0.007 | 0.865 | 0.795 | 0.811 | 0.801 | 0.807 | 0.796 |
| 0.01 | 0.821 | 0.792 | 0.798 | 0.800 | 0.803 | 0.768 |
| 0.03 | 0.791 | 0.724 | 0.771 | 0.657 | 0.694 | 0.674 |

The dimension of concatenated node features is a key factor for model performance. We conduct the cross-validation to assess the impact of node dimensions (Table 3). It indicates that the performance of the model becomes better when the dimensionality of the node increases from 32 to 128. The model achieves best performance when the node dimension is 128. However, as the node dimension continues to increase, the performance of the model decreases. The default value of node dimension is set to 128.

**Table 3.** The impact of node dimensions on model performance

| Node dim | AUC | AUPR | Acc | Pre | Recall | F1-Score |
| --- | --- | --- | --- | --- | --- | --- |
| 32 | 0.873 | 0.775 | 0.787 | 0.761 | 0.784 | 0.759 |
| 64 | 0.879 | 0.798 | 0.813 | 0.795 | 0.802 | 0.792 |
| **128** | **0.886** | **0.817** | **0.824** | **0.808** | **0.817** | **0.804** |
| 160 | 0.882 | 0.796 | 0.814 | 0.797 | 0.808 | 0.788 |
| 192 | 0.865 | 0.761 | 0.776 | 0.758 | 0.765 | 0.762 |

### 3.3 Changes of attention scores

We use attention mechanism to determine the influence of each type of edge. During the training, changes in the weights of each type of edge reflect the importance of the pathogenic process (Fig. 2). CM stands for circRNA-miRNA matrix. CD stands for circRNA-disease Matrix. MD stands for miRNA-disease Matrix. I is an abbreviation for identity matrix. The CD matrix has the highest allocated weight, implying that it has the greatest influence on the final prediction accuracy. CM and MD are also important for accuracy. I has minimal impact on accuracy. The conclusion is consistent with the actual biological situation.

**Fig. 2.**
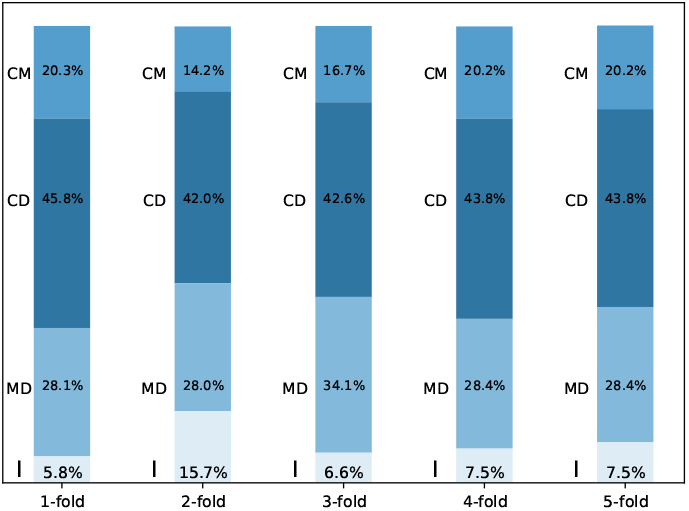
Changes of attention values during 5-fold cross-validation

### 3.4 Comparison with other methods

We contrast CDHGNN with five state-of-the-art methods, including KGA-NCDA (Lan *etal*., 2021), MGRCDA (Wang *et al*., 2021), CDASOR (Lu *et al*., 2020), GCNCDA (Wang *et al*., 2020) and NSL2CD (Xiao *et al*., 2021). We also evaluate the effects of edge-weighted graph attention network and heterogeneous graph transformer network separately. The first model composed of un-weighted GAT and heterogeneous GTN is marked as CDHGNN_u_. The second model composed of edge-weighted GAT and homogeneous neural network is marked as CDHGNN_h_. The detailed comparison results are shown in Table 4. We can see that CDHGNN achieves the best performance (AUC:0.886, AUPR:0.817, Accuracy:0.824, Precision:0.808, Recall:0.814, F1-score:0.804). The edge-weighted GAT improves the performance (AUC:1%, AUPR:2%, Acc:2%, Pre:1%, Recall:1%, F1-score:0.7%). The heterogeneous neural networks improves even more (AUC:1.9%, AUPR:4%, Acc:4%, Pre:3.3%, Recall:1.6%, F1-score:0.9%). The detailed AUC results are shown in Fig. 3. The auc value of CDHGNN is much higher than that of other methods, indicating that the accuracy of CDHGNN is higher. In addition, we compare the percentage of correctly retrieved associations from top 10 to top 40 predictions (Fig. 4). All the results show that not only edge-weighted GAT effectively improve the model, but also heterogeneous graph neural networks make the model more accurate. Compared with edge-weighted GAT, the utilization of a heterogeneous graph neural network improves the accuracy more significantly. It indicates that the introduction of multi-source biological information can enhance prediction.

**Table 4.**
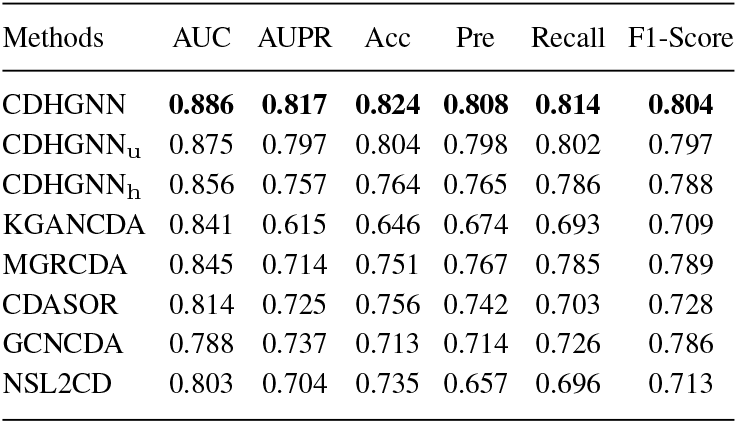
The performance comparison with state-of-the-art methods.

**Fig. 3.**
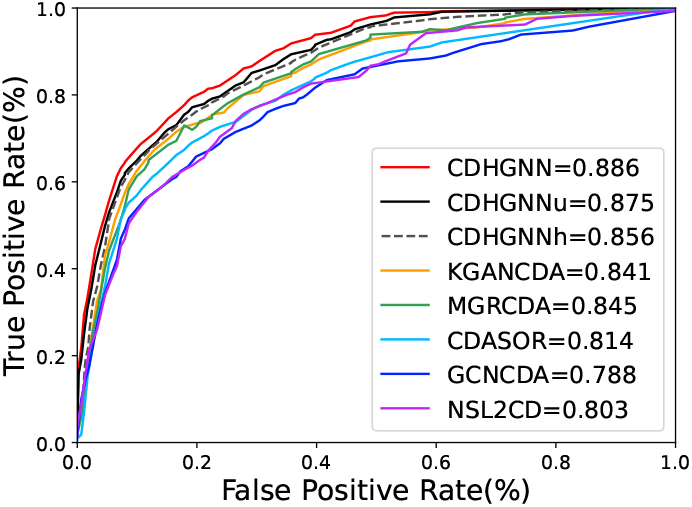
The performance comparison with state-of-the-art methods in terms of AUC values.

**Fig. 4.**
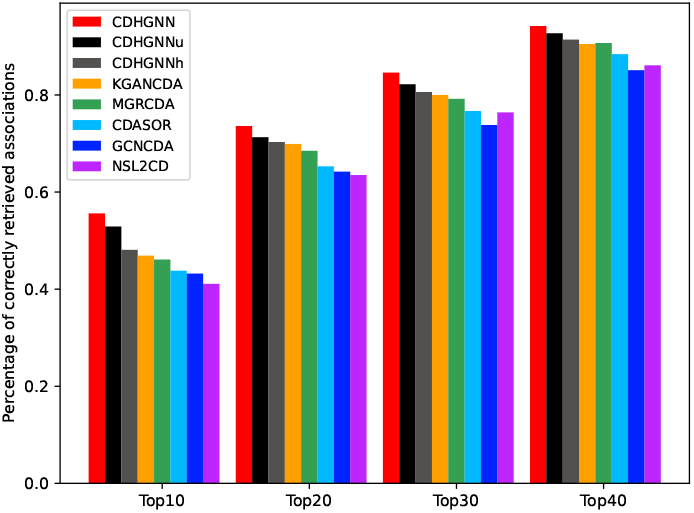
Percentage of correctly retrieved associations

### 3.5 Case studies

Case studies are conducted to assess the prediction capacity with existing literature and public databases. Sort predicted relationships in descending order after training the model with all the empirically verified circRNA-disease associations. Table 5 shows 14 of the top 20 predicted connections were verified. Originated from vacuolar ATPse assembly factor, hsa_circ_0091702 mitigates sepsis-correlated acute kidney injury by regulating miR-9-3p/SMG1/inflammation and oxidative stress (Shi *et al*., 2020). Circ-AKT3 (hsa_circ_0000199) is related to acute kidney injury via miR-144-5p/Wnt/*β*-catenin pathway (Xu *et al*., 2020). CircFUT8 (hsa_circ_0003028) sponges miR-570-3p and regulate the miR-570-3p/KLF10 axis as a tumor suppressor in bladder cancer (He *et al*., 2020). Binding with miR-296-5p, hsa_circ_0000515 activates the cell growth of bladder cancer (Cai *et al*., 2020). Fastening miR-200a-3p, exosomes-mediated transfer of circ_UBE2D2 (hsa_circ_0005728) enhances tamoxifen resistance in breast cancer (Hu *et al*., 2020). Sponging miR-532-3p, circRNA_103809 (hsa_circ_0072088) represses cell proliferation and metastasis of breast cancer (Liu *et al*., 2020). Participating in the miR-135a-5p/EMT axis, circRNA_0001946 (hsa_circ_0001946) functions as a tumor promoter (Zeng *et al*., 2020). Inhibiting colorectal cancer cell proliferation by its knockdown and regulating miR-296-5p/RUNX1 axis, circ_0000512 (hsa_circ_0000512) is a promising therapeutic target for colorectal cancer (Wang *et al*., 2020). Circ-RanGAP1 (hsa_circ_0063526) promote gastric cell progression by mediating miR-877-3p/VEGFA axis (Lu *et al*., 2020). In gastric cancer, hsa_circ_0004872 related to a negative regulatory loop hsa_circ_0004872/miR-224/Smad44/ADAR1 functions as a tumor suppressor (Ma *et al*., 2020). Acting as endogenous RNA for miR-654-3p, circRHOBTB3 (hsa_circ_0006404) inhibits cell growth of gastric cancer by promoting p21 signaling pathway (Ma *et al*., 2020). Insulating miR-17, circ-ITCH (hsa_circ_0001141) functions as a tumor-suppressor factor by the Wnt/*β*-catenin signaling pathway in gastric cancer (Peng *et al*., 2020). Sponging miR-134-5p which activates BTG-2 expression, circZNF609 (hsa_circ_0000615) represses proliferation and migration of glioma cell (Tong *et al*., 2020). Targeting the miR-520a-5p/CDK4 regulatory axis, exosome-transmitted hsa_circ_0014235 activates malignant development of non-small cell lung cancer (Xu *et al*., 2020). The details of validated pathways for the top-20 predicted circRNA-disease associations is shown in Fig. 5. MiRNAs linked to circRNAs and involved in diseases have been discovered, which could aid in the discovery of new pathways.

**Table 5.**
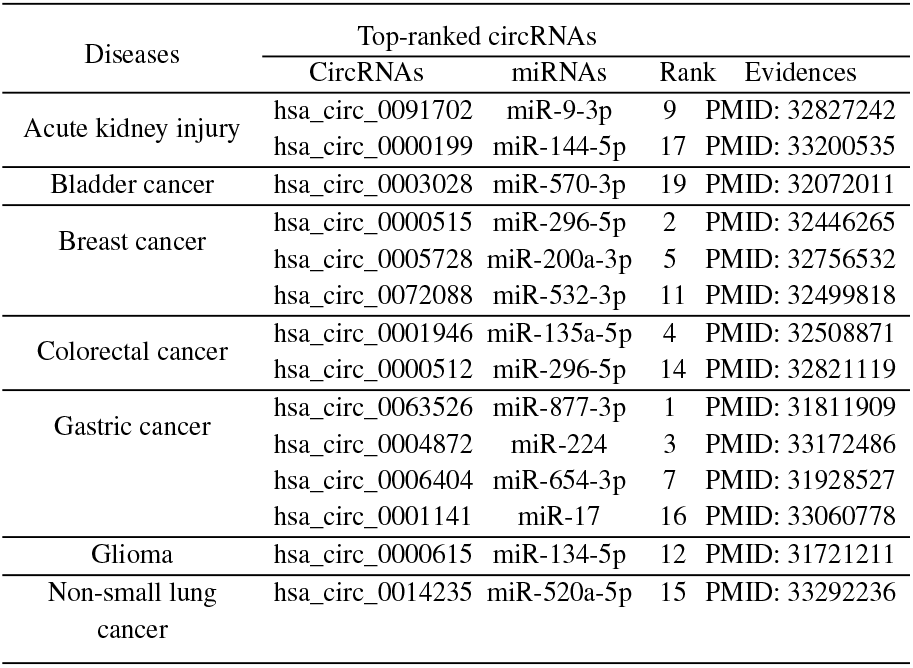
Validation of top-20 rank predicted associations.

**Fig. 5.**
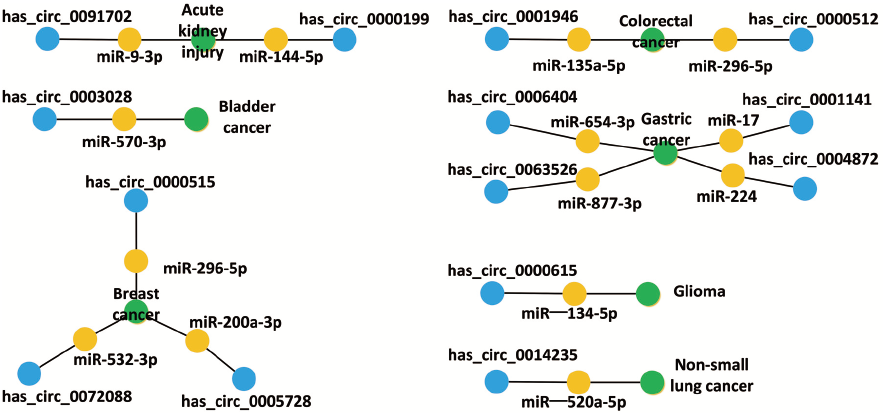
The validated pathways for the top-20 predicted circRNA-disease associations.

## 4 Conclusion

CircRNAs and illnesses are linked by their range of biological roles. Meanwhile, the special structure makes it a promising biomarker for the treatment of diseases. However, existing methods lack consideration of multi-source data heterogeneity. In this work, we devise a model to predict possible circRNA-disease associations based on a heterogeneous graph neural network. We created a unique edge-weighted graph attention network to grasp node features since edge weights convey the relevance of associations between nodes. We adopt an attention mechanism to learn contextual information and assign attention scores on the meta-path to infer potential associations. The experimental results reveal that CDHGNN outperforms state-of-the-art methods with comparable accuracy. It is worth noting that CDHGNN can find molecular connections and the relevant pathways in pathogenesis.

Although CDHGNN performs admirably in terms of predicting probable associations, it still has several flaws that need be investigated further. For example, the pathogenic process of disease is a very complex molecular activity process. CDHGNN uses biological information among circRNA, miRNA and disease. It would be preferable if additional relevant biomolecular data were integrated to train the model. Furthermore, the molecular structure, which provides unique information about biomolecules, may be enhanced by adding more accurate features to the prediction model.

